# GUANACO: A Unified Web-Based Platform for Single-Cell Multi-Omics Data Visualization

**DOI:** 10.1101/2025.09.18.677070

**Authors:** Xiyuan Zhang, Marian Kuddus, Qianqian Xia, Yizhou Hu, Ping Chen

**Author notes:** Correspondence: Yizhou Hu; Ping Chen.

## Abstract

While single-cell multi-omics has advanced our understanding of cellular heterogeneity, the complexity of these datasets remains a barrier for non-computational users. Here, we introduce GUANACO, a Dash-based Python package for interactive, code-free visualization of single-cell RNA-seq and ATAC-seq data. GUANACO integrates matrix- and genome-track views, supporting flexible cell/gene selection, statistical testing, and transcription factor binding site exploration. Its user-friendly interface offers colorblind-friendly palettes, intuitive controls, and options for generating publication-ready figures. With a low memory requirement and cost-effective deployment, GUANACO facilitates seamless sharing and reproducible research, empowering researchers to explore, interpret, and communicate single-cell insights without extra coding.

## Background

Advances in single-cell technologies have substantially improved the resolution at which cellular heterogeneity and dynamics can be studied [1]. As a result, vast amounts of single-cell datasets have been generated and made publicly available, providing valuable resources for developing new biological hypotheses. However, the scale and high dimensionality of these data can be overwhelming for experimental biologists, particularly those without computational expertise [2]. This challenge becomes even more pronounced in multi-omics studies, where different molecular layers have to be linked for biological interpretation, highlighting the pressing need for intuitive, user-friendly visualization tools that can help the broader research community translate processed data into meaningful biological insights.

Visual exploration of processed single-cell data in coding environments such as Jupyter Notebook typically begins with AnnData [3] or Seurat [4] objects generated from preprocessing pipelines, followed by iterative inspections of gene expression and metadata (e.g., cell type, individual, and disease state) [5]. This process is often repetitive, requiring frequent script modifications to create alternative plots, which can make it tedious, error-prone, and particularly challenging when handling large datasets or testing multiple gene-metadata combinations. In addition, refining figures for presentations and publications in such environments usually involves code adjustments to fine-tune visual parameters and aesthetics [6]. While coding environments provide flexibility for data analysis and custom visualization, they do not offer a straightforward or easily shareable solution for broad biological exploration, particularly for researchers without coding expertise.

To address these challenges, web-based visualization platforms such as CELLxGENE [7], and the UCSC Cell Browser [8] have been developed, offering intuitive graphical user interfaces that can greatly facilitate data exploration and dissemination. Nevertheless, their utility for routine visual inspection could be further improved by expanding plot libraries and interactive controls, enabling biologists to navigate and interpret data more efficiently. More recently, tools like Vitessce have emerged for multimodal single-cell data visualization [9]. While powerful, Vitessce often requires computational skills to configure and optimize visualizations. Although these platforms represent valuable resources, each serving different purposes, there remains a need for user-friendly frameworks that integrates built-in statistical functionality, expanded plotting options, and more flexible dashboard layouts to streamline exploratory analysis. Shinycell2 [10] addresses part of this need by offering configurable graphical settings, publication-quality static figures and basic statistical tests. However, its static outputs come at the cost of interactivity, and features such as dynamic filtering or on-the-fly parameter tuning remain promising areas for development. The growing complexity of single-cell datasets calls for adaptable and interactive visualization frameworks that can extend functionality to address emerging needs and, in turn, accelerate data exploration and biological discovery.

Here, we introduce **GUANACO** (**G**raphical **U**nified **A**nalysis and Navigation of **C**ellular **O**mics), a Python-based platform designed for biologists to explore processed single-cell multiomics data (scRNA-seq and scATAC-seq), with support for both AnnData [3] and MuData [11] input formats. GUANACO combines adaptable and interactive data exploration with publication-ready visualizations and facilitates seamless sharing of fully integrated, interactive webpages. With an emphasis on human-centred design and flexible, purpose-driven workflows, it offers an accessible and lightweight solution that integrates smoothly into existing preprocessing pipelines while minimizing the manual effort required for visual inspection.

## Results

### GUANACO streamlines multimodal single-cell data exploration and presentation through a unified web interface

GUANACO supports integrated visualization of scRNA-seq and/or scATAC-seq data from single or multiple studies through a unified, browser-based interface. Each study is presented in a dedicated tab comprising two views: a matrix-based single-cell omics view (AnnData/MuData) and a coordinate-based genome track view, allowing seamless exploration of gene expression and chromatin accessibility landscapes. GUANACO can be launched with a single-line command pointing to a configuration *JSON* file, requiring only the *JSON* filename in its minimal form, with additional customizations optional. Cloud deployment for data sharing is also straightforward, requiring minimal coding effort (Fig. 1a).

**Figure 1.**
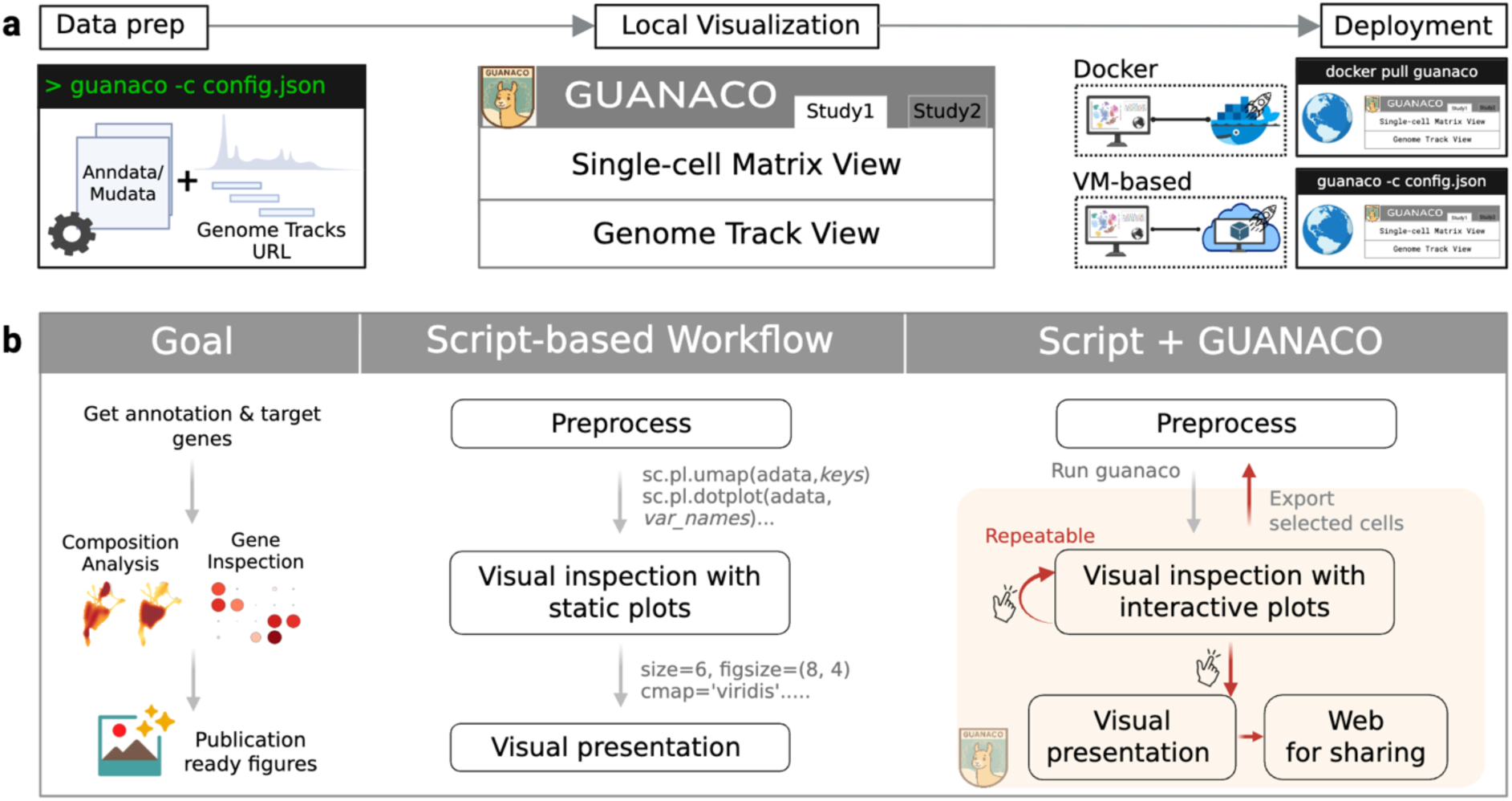
The overview of GUANACO. (a) GUANACO workflow. Left: JSON configuration setup combining AnnData/MuData objects with optional genome track URLs. Middle: Dual-view interface for Single-cell Matrix View and Genome Track View, supporting multiple studies in a tabbed interface. Right: GUANACO offers flexible deployment options through Docker container or system’s default Python environment. (b) Comparison of traditional script-based versus GUANACO workflows for single-cell data visualization. Script based workflow (left) requires repetitive coding for visual inspection with static plots (e.g., ‘sc.pl.umap’, ‘sc.pl.dotplot’ in Scanpy [12] with manual parameter adjustments), followed by additional code modifications for presentation-ready figures. The GUANACO workflow (right) streamlines this process: after preprocessing, a single command launches an interactive, shareable interface for intuitive data exploration and figure creation with mouse clicks. Selected cells can be exported back to the preprocessing pipeline for subcluster analysis and subsequently visualized in GUANACO with updated annotations (bidirectional workflow).

Figure 1b illustrates how GUANACO streamlines the traditional single-cell analysis workflow. Instead of writing repetitive code for each plot, users can launch a suite of synchronized and reusable interactive visualizations directly from preprocessed data. Importantly, selected cell populations can be propagated across all visualizations and fed back into the preprocessing pipeline, making visualization an integral part of the discovery process. This approach aligns with current single-cell analysis practices, where visualization actively drives discovery. In addition to supporting exploration, GUANACO enables the creation of publication-quality figures through intuitive graphical controls, allowing users to adjust colors, sizes, and layouts without writing code. Users can also export static figures(.svg) or share interactive visualizations through a web link.

To illustrate GUANACO’s interface, we use integrated single-cell multi-omics data from two peripheral blood mononuclear cell (PBMC) datasets (10x10k and 10x3k; see Methods). The top panel of the interface displays a data summary, including dataset descriptions provided in the configuration file and dataset dimensions. Data selection for matrix-based data visualization is structured into five hierarchical levels (highlighted in red in Fig. 2a). Level 1 (Modality Selection):

**Figure 2.**
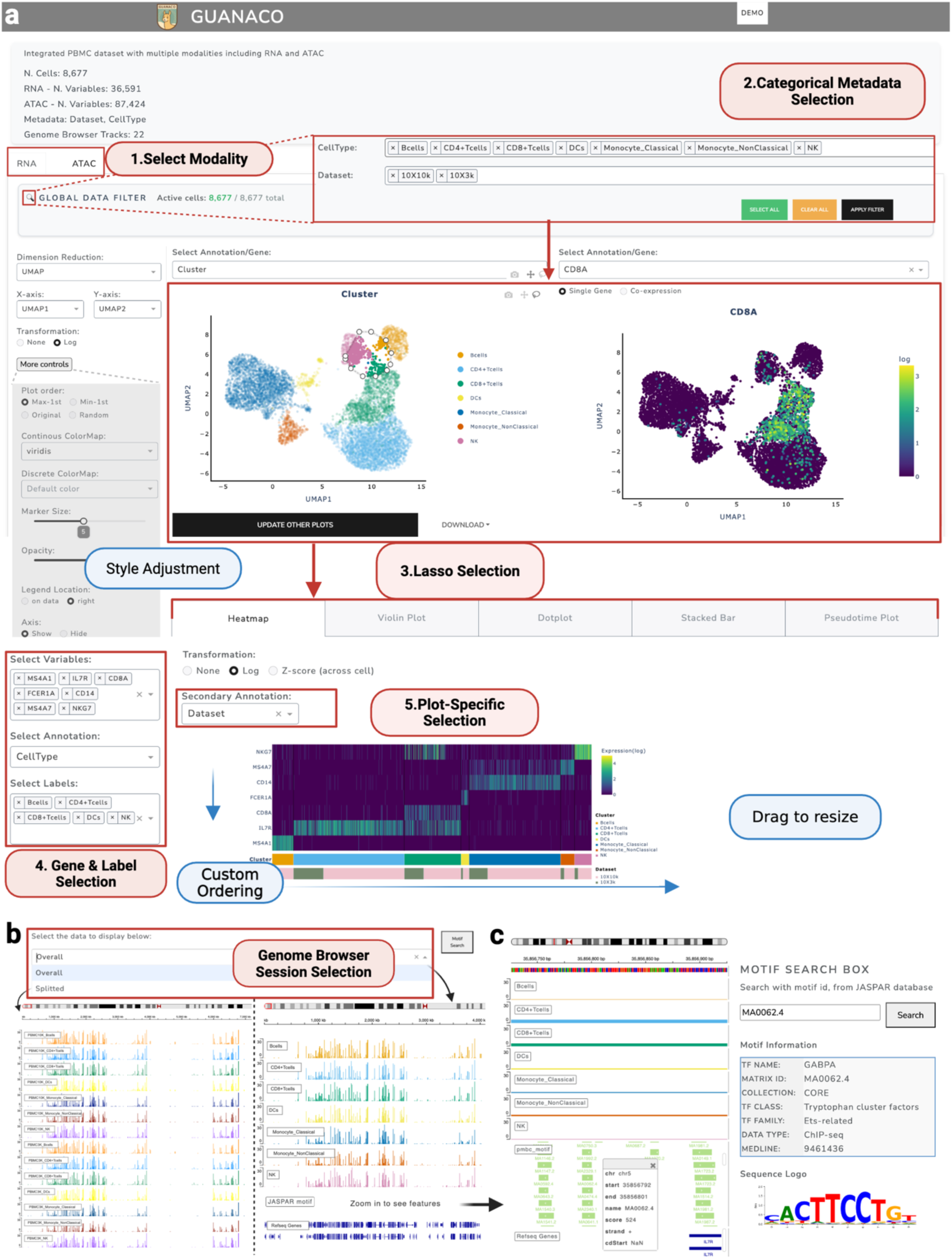
GUANACO user interface for matrix-based and genome track visualizations. (a) Matrix-based visualization interface. Red highlights indicate hierarchical data selection steps, including modality selection, global metadata filters, embedding-based lasso selection, gene and label specification, and plot-level customization. Blue highlights indicate visual adjustments such as style configuration, ordering, and resizing. (b) Genome track interface using IGV integration [14]. Two viewing modes are available: ‘Overall,’ which shows peaks grouped by cell type, and ‘Split,’ which separates peaks by both dataset and cell type. Red highlights show session selection options. (c) Predicted transcription factor binding motifs in the *IL7R* promoter region with cell type-specific chromatin accessibility. The validated GABPA binding motif (MA0062.4) is highlighted, with additional information displayed in the motif search box.

For multi-omics datasets, users first select the modality tab (e.g., RNA or ATAC); Level 2 (Global Selection): Users can then filter cells globally using categorical metadata, with full flexibility to combine or exclude groups across all variables; Level 3 (Lasso Selection): This view displays the globally filtered cells in a dimensional-reduction plot (e.g., UMAP) with interactive lasso selection, designed for bench scientists to directly explore cellular heterogeneity within any cell population of interest. After clicking Update Plots, selected cells are passed to all downstream visualizations, enabling focused, code-free exploration of their molecular profiles; Level 4 (Gene and Label Selection): Within the selected subset after global and lasso selections, users can specify genes and annotations to construct plots such as heatmaps, violin plots, and dot plots; Level 5 (Plot-Specific Selection): Each plot type includes its own set of configurable options, such as secondary annotations in heatmaps, toggling boxplots in violin plots, or hierarchical clustering in dot plots. Moreover, GUANACO offers additional graphic setting features highlighted in blue (Fig. 2a), including a global style control panel for adjusting plot aesthetics (e.g., colormap settings), drag-to-resize functionality for resizing plots, and custom row/column ordering based on the order of input selections, which can be fully rearranged via the left-side controls.

For genome track visualizations, users can view ATAC peaks grouped by user-defined criteria in different track sessions. For example, in Fig. 2b, the split-session view shows peaks stratified by both cell type and dataset, whereas the overall-session view groups peaks by cell type alone. When zooming into specific peak regions, the motif track displays transcription factor binding motifs from the JASPAR CORE database. For instance, Fig. 2c illustrates a zoomed-in view of the *IL7R* promoter region (−100 bp from the TSS), a gene highly expressed in T cells and encoding the interleukin-7 receptor essential for T cell development [13]. Our genome track view reveals increased chromatin accessibility at this promoter in both CD4+ and CD8+ T cells, overlapping with a GABPA binding motif (MA0062.4). This finding is consistent with previous evidence that GABPA, an ETS-related transcription factor, positively regulates *IL7R* transcription and plays a critical role in lymphocyte development and survival [13].

### Gallery of GUANACO visualizations

GUANACO delivers a comprehensive suite of visualization modules designed for single-cell data exploration in both matrix-based and coordinate-based genome track views. These modules facilitate downstream exploration, such as inspecting cell state differences across target groups, tracing transitions along trajectory paths, and deriving mechanistic insights through multimodal comparisons. The interface seamlessly translates user inputs, such as gene/annotation selection, cell selection, and visualization settings, into interactive visual outputs via intuitive controls, including dropdown menus, sliders, and plot type selectors (Fig. 3a). Core visualization modules for single-cell matrix data are summarized in Fig. 3b and as below:

1. Dimensionality reduction: Interactive scatter plots (e.g., UMAP, t-SNE, PCA) are displayed side by side with synchronized zooming. Each plot supports visualization of both gene expression and metadata, as well as co-expression analysis with adjustable expression thresholds. Lasso selection can be used to highlight specific cell populations, and selected cells are propagated across all visualizations. Real-time controls adjust visual parameters (color, point size, opacity), supporting both categorical annotations and continuous gradients for in-depth cell state exploration.
2. Heatmap: GUANACO generates single-cell level heatmaps with support for up to two annotation layers, which can include categorical metadata or continuous variables such as pseudotime. The second layer can be used to highlight batch effects, subclusters, disease conditions, or reveal temporal patterns by ordering cells along pseudotime. In addition, cluster boundaries are drawn to improve accessibility.
3. Violin plot: GUANACO provides stacked violin plots to display the expression levels of multiple genes in a compact view, as well as split or grouped violin plots with statistical information for individual genes, with optional overlays of boxplots and individual data points. In the split and grouped violin view, four analysis modes with selectable statistical tests are available. *Mode 1* compares a single metadata variable (Mann–Whitney U or t-test for two groups; Kruskal– Wallis or ANOVA for more than two groups). *Mode 2* performs faceted comparisons within subgroups. *Mode 3* applies linear models with confounders (e.g., expression ∼ obs1+ obs2), such as cell type and batch. *Mode 4* uses mixed-effects models to account for hierarchical data structures (e.g., expression ∼ obs1+ (1 | obs2)), such as condition and patient, where patients are nested within conditions.
4. Dot plot: The dot plot summarizes expression data at the cluster level, with color intensity representing mean expression and dot size reflecting the fraction of expressing cells, while optional hierarchical clustering (rows, columns, or both) with configurable linkage and distance metrics can reorder the axes. GUANACO extends traditional dot plots by automatically scaling dot sizes to improve the visibility of genes with low expression proportions and by offering standardization options (by gene or by cluster) to enhance visual contrast. A matrix view is available as an alternative for cases when expression patterns are prioritized over cell fractions.
5. Stacked bar plot: Summarizes cell type compositions based on user-defined metadata (e.g., dataset). Bars along the x-axis can be reordered either by adjusting their input order or via drag-and-drop. Both proportional and absolute counts are supported. When only cell type is selected, standard bar plots are displayed to show either the proportion or absolute counts of each cell type across all cells.
6. Pseudotime gene plot: GUANACO also provides pseudotime gene plots when the input object includes pseudotime metadata. This visualization displays gene expression (Y-axis) along pseudotime (X-axis) as a scatter plot with annotation-based coloring, while a smoothed trend line fitted by polynomial regression illustrates the expression trajectory.

**Figure 3.**
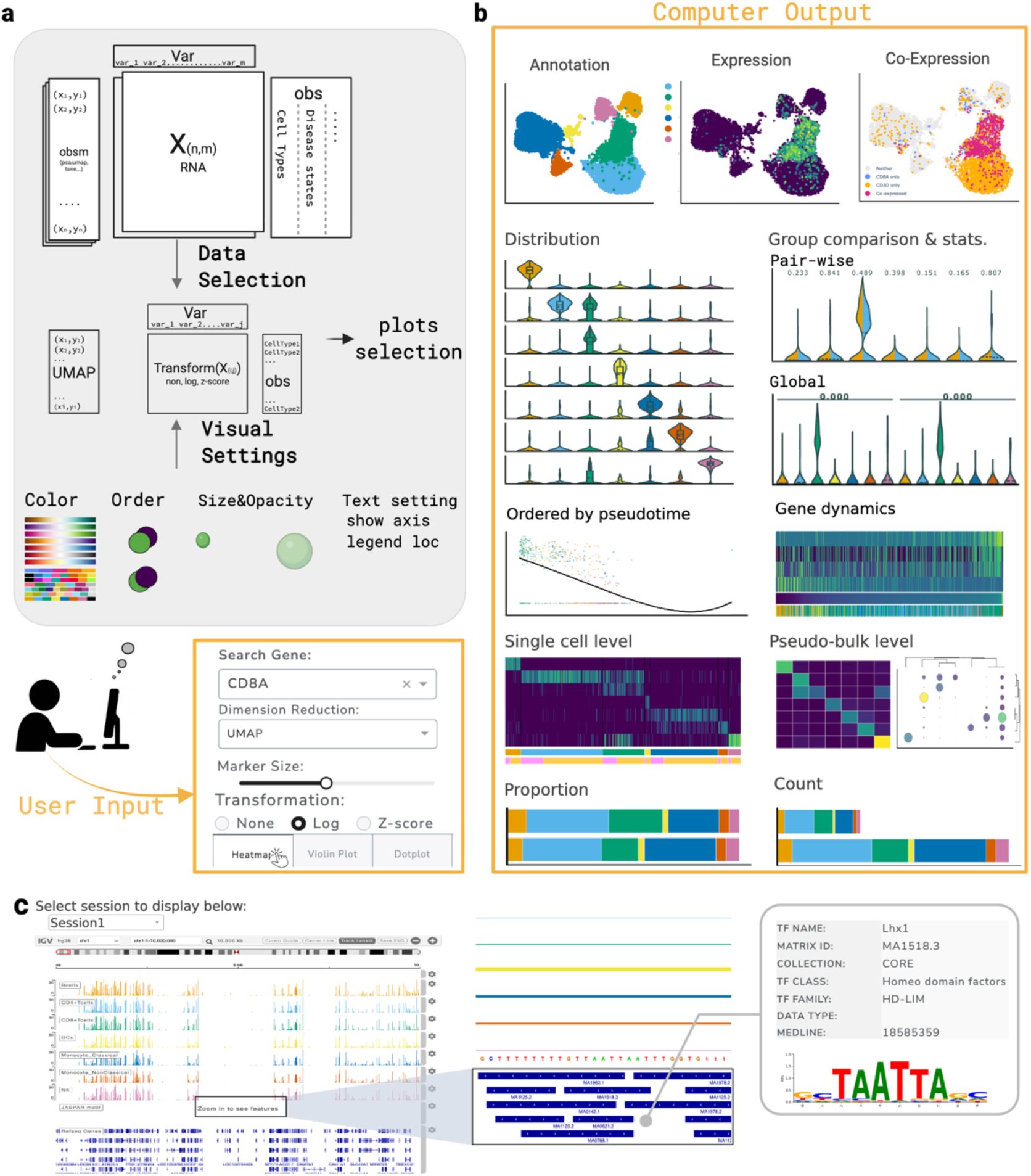
Gallery of plot types in GUANACO. (a) GUANACO control interface for translating user inputs into visual outputs. Data selection allows choosing among multiple dimensionality reductions and subsetting observations and variables from an AnnData object. Visual settings provide fine control over color palettes, point ordering, size, opacity, and text display. Inputs are captured through search boxes, dropdown menus, sliders, and radio buttons, enabling code-free exploration. (b) Single-cell matrix data visualization. Available plot types include side-by-side dimensionality reduction plots, violin plots for distributions (grouped and split styles with statistical testing), heatmaps at single-cell and pseudo-bulk levels, dot plots, and stacked bar plots for proportions and absolute counts. (c) Genome track visualizations. GUANACO supports multi-session coordinate-based data types such as ATAC-seq peaks, motifs, and SNPs. Motif details can be explored through a built-in searchable database.

For coordinate-based genome track data, GUANACO integrates an IGV-based genome browser optimized for single-cell multi-omics exploration (Fig. 3c). Tracks are color-synchronized to reflect annotations from the AnnData object, ensuring consistent interpretation across modalities. Users can switch between predefined sessions (e.g., by cell type or condition) using dropdown menus, with each session retaining full IGV functionality, including genome navigation, track customization, and support for standard formats (e.g., BED, BigWig, VCF). A built-in motif explorer allows direct querying of motif identifiers from the JASPAR database [15], returning associated transcription factors alongside detailed descriptions and motif sequence logos.

### Flexible, intuitive, and accessible interactive visualization

GUANACO enhances single-cell data exploration with a responsive and intuitive interaction interface. Its interactive features transform traditional dimensional reduction plots into powerful selectors, supporting real-time updates to downstream visualizations (heatmaps, violin plots, dot plots, and bar plots) and enabling synchronized side-by-side exploration through linked zooming. Selected cells can be exported as a cell_id text file for in-depth analysis with user-preferred pipelines (Fig. 4a). Interactive tooltips reveal precise expression values on hover while keeping a clean visual layout. This is particularly valuable in dot plots, where dot size alone may give an imprecise sense of magnitude (Fig. 4b).

**Figure 4.**
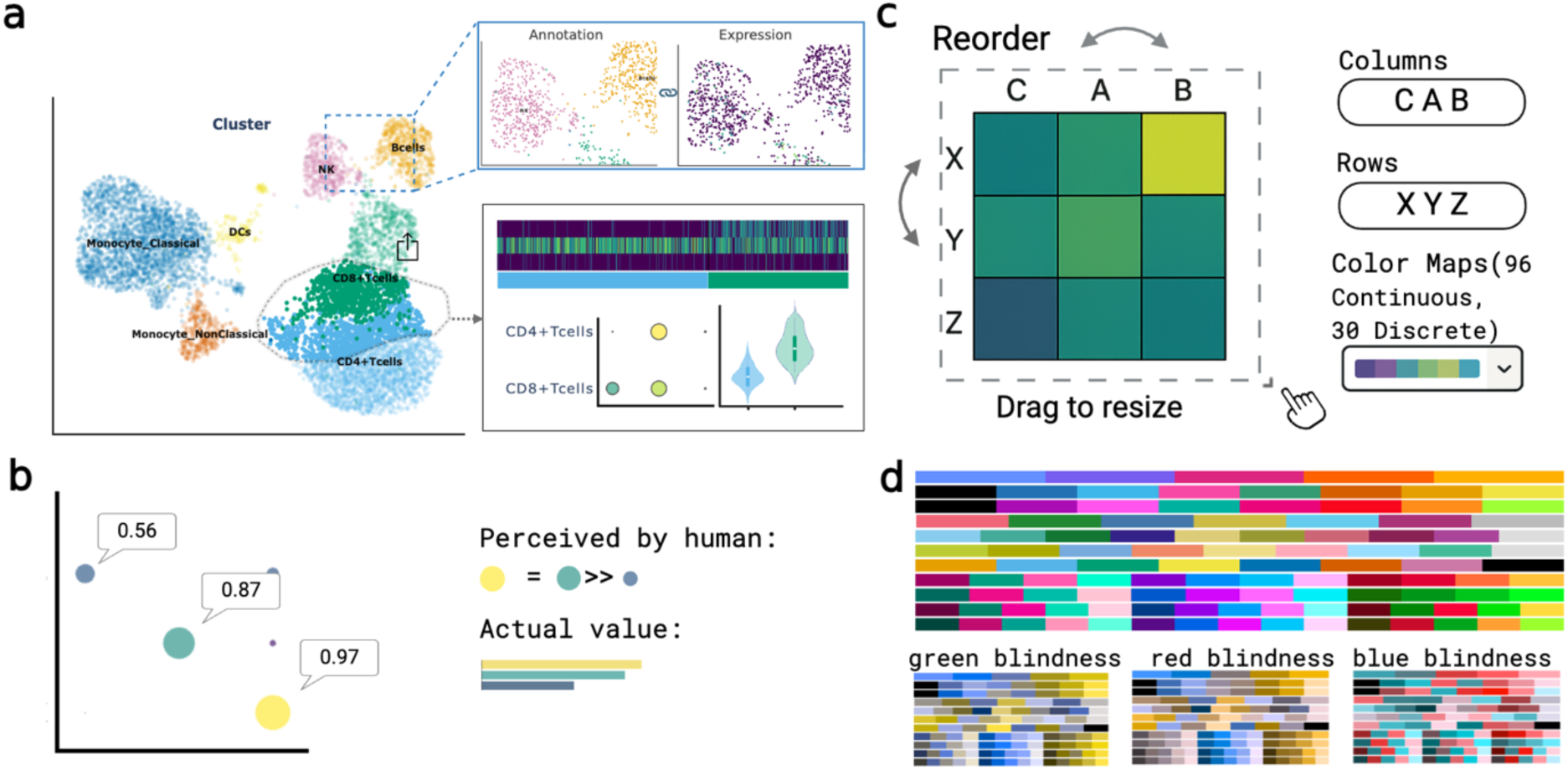
Flexible, intuitive and accessible interactive features in GUANACO. (a) Dimensional reduction plots support zooming and synchronized views between gene expression and annotation panels. Cells selected via lasso can be propagated to other visualizations (e.g., heatmap, violin plot, dot plot) and exported for in-depth analysis. (b) Tooltip overlays display exact expression values on hover, helping to address perceptual biases in dot plots. (c) Multi-feature plots allow interactive reordering of rows and columns, resizing by drag and drop, and flexible color selection via dropdown menus, offering 96 continuous and 30 discrete colormaps. (d) Eleven newly developed colorblind-friendly colormaps in GUANACO, with simulated views generated by Coblis [18] for three types of color vision deficiency at the bottom.

For figure generation, GUANACO prioritizes direct manipulation and visual flexibility. Users can resize plots simply by dragging and reorder features by adjusting the input data order (Fig. 4c), which provides a more intuitive and faster approach than entering numeric values or manually reorganizing data. A broad range of visual customization options are available, including an extensive color map library with 96 continuous and 30 discrete colormaps. Notably, 11 of the discrete colormaps are newly developed colorblind-friendly palettes in GUANACO (Fig. 4d), designed in accordance with established guidelines for colorblind-accessible figure creation [16, 17].

### Comparative evaluation of GUANACO against existing single-cell visualization platforms

Table 1 compares the key features of GUANACO with other state-of-the-art single-cell visualization tools, focusing on both data exploration and presentation capabilities (detailed comparison in Additional file 2). Client-side tools such as CELLxGENE, UCSC Cell Browser, and Vitessce deliver scalable browsing and effortless data sharing, making them well suited for initial exploration and public dissemination. Their natural next step lies in embedding statistical assessments for group-wise comparisons and a broad range of plot types to further streamline data exploration and biological hypothesis generation. Moreover, enhanced and flexible graphical customization would allow users to generate publication-ready figures directly, without the need for additional figure polishing. ShinyCell2 offers a user-friendly interface designed for biologists, though its exploratory capacity could be further expanded with features such as interactive subsetting and multifactor comparisons. GUANACO positions itself as a complementary solution that unites these strengths while addressing emerging needs. It bridges current gaps by supporting both flexible, interactive data exploration and high-quality presentation across datasets of typical single-cell scale. By integrating a range of interactive features while maintaining a low memory footprint, GUANACO ensures a seamless transition from exploration to publication-ready output.

**Table 1.**
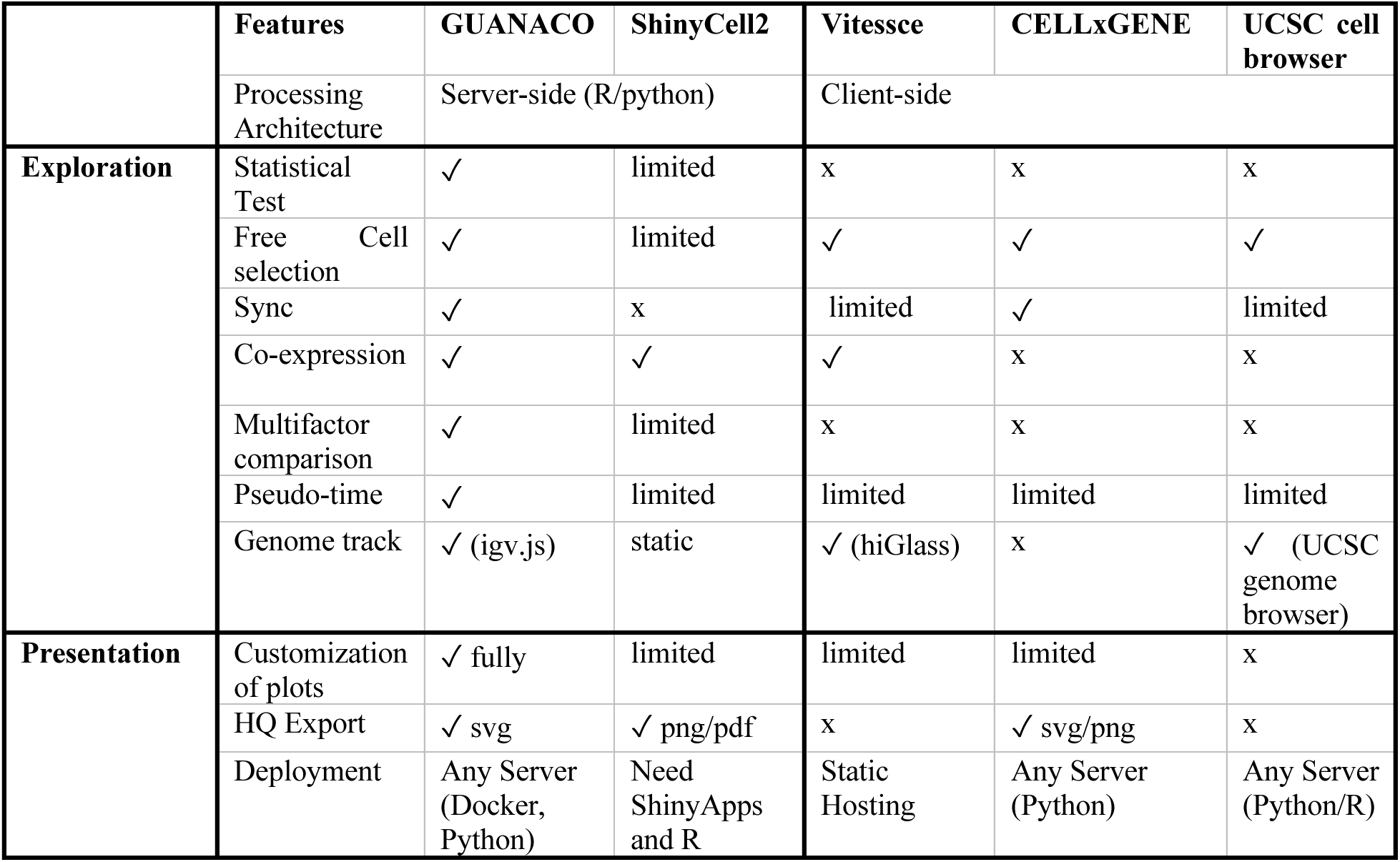
Comparison of existing tools with GUANACO in terms of data exploration and presentation. Checkmarks (✓) indicate support, “x” indicates no support, and “limited” denotes restricted implementations of a feature (e.g., only basic statistical functions, categorical-only selection, or support for a subset of methods).

To evaluate how well each tool supports common biological tasks, we conducted a task-driven assessments. First, we examined subcluster identification using the same PBMC datasets shown in Fig. 2. Taking the NK cell marker *NKG7* as an example, we observed high *NKG7* expression in a subset of CD8+ T cells (Fig. 5a), suggesting the presence of an NKT cell subset or potential NK-like cytotoxic phenotype [19] not captured by the original annotation. With GUANACO, users can interactively subset CD8+ T cells, filter them by metadata variables (e.g., condition and batch), or zoom into the NKG7-high region to compare differences across defined groups. For example, in GUANACO lasso-selected cells as shown in Fig. 5a, cell composition across dataset batches is visualized with stacked bar plots, while *NKG7* expression levels are compared between datasets using split violin plots with associated p-values. These views allow users to assess whether the observed differences reflect batch effects or a true CD8⁺ T cell subcluster. Selected cells can also be exported for further investigation using user-preferred pipelines. By comparison, ShinyCell2 only supports filtering by a single metadata variable at a time, making conditional comparisons within each cell type infeasible. Furthermore, since ShinyCell2 only provides static plots, users cannot interactively select specific cells or regions of interest from embeddings for further analysis. Vitessce supports lasso selection but lacks dynamic linkage to metadata filtering, requiring prior code-based subsetting for cross-condition comparisons.

**Figure 5.**
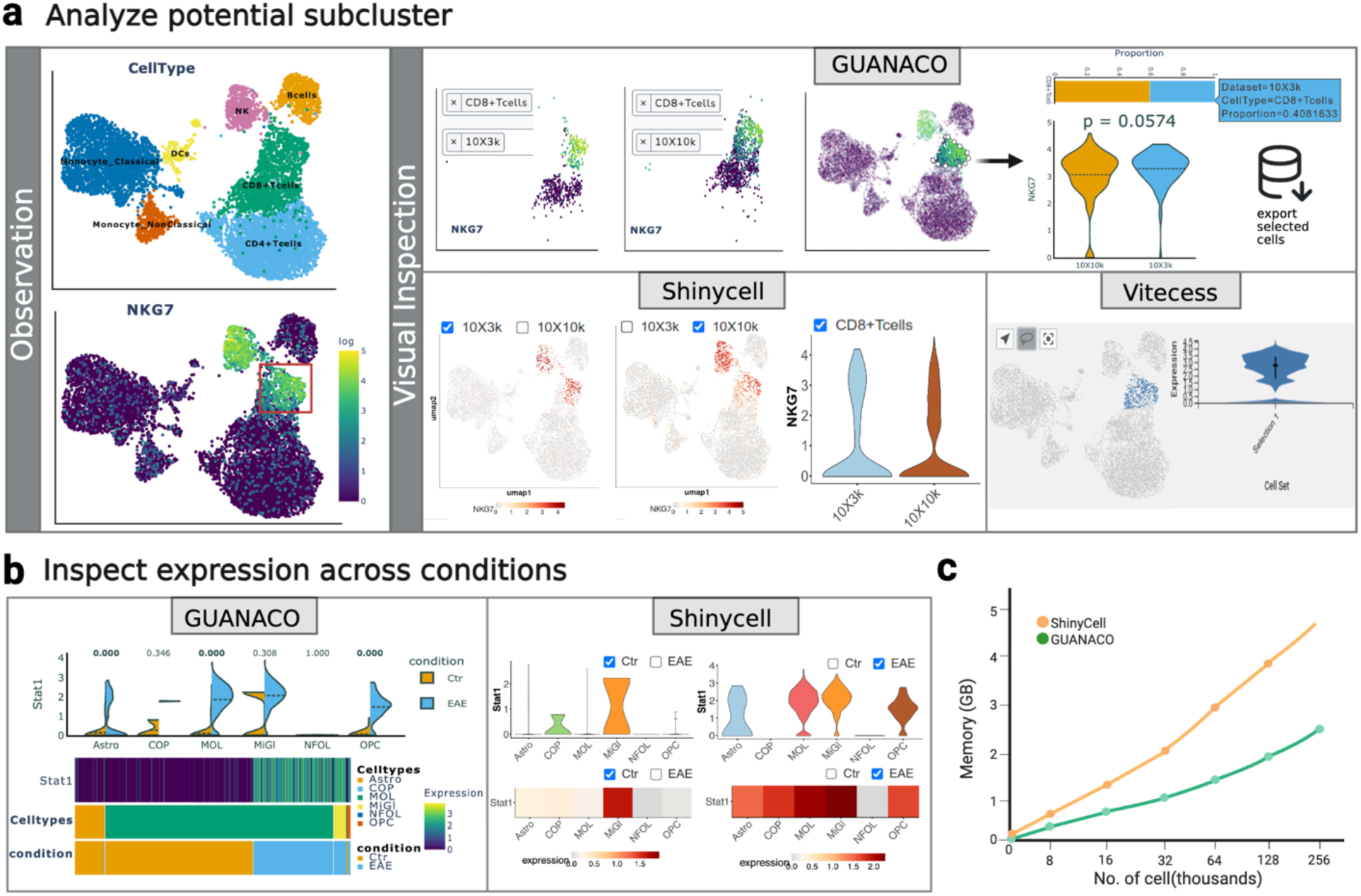
Detailed visual and performance comparison of GUANACO and other single-cell visualization tools. (a) Left: Initial observation using UMAP plots colored by cell type and *NKG7* expression reveals elevated *NKG7* signal in a subset of CD8⁺ T cells. Right: GUANACO enables interactive selection of CD8⁺ T cells stratified by dataset sources (10X3k and 10X10k). The *NKG7*⁺ region can be zoomed in for further inspection. A bar plot displays the sample composition of the selected region, while split violin plots compare *NKG7* expression in the selected region between datasets (10X3k vs 10X10k), annotated with statistical significance. An export icon indicates that selected cells can be saved for further analysis using user-preferred pipelines. ShinyCell2 shows *NKG7* expression across datasets with a single metadata filter (CD8⁺ T cells), but only static violin plots are available, and region-specific selection is not supported. Vitessce allows lasso selection in UMAP space, but selected cells are not dynamically linked to metadata filters, limiting condition-specific comparisons. (b) Cross-condition gene expression inspection in a published oligodendroglia dataset comparing healthy (Ctr) and EAE mouse samples. Left (GUANACO): Split violin plots show *Stat1* expression across cell types, with integrated p-values indicating significant differences in Astro, MOL, and OPC populations. The heatmap below visualizes single-cell *Stat1* expression across both cell types and conditions, annotated by cell identity and condition. Right (ShinyCell2): Violin plots and heatmaps are grouped by condition, requiring manual switching to inspect comparisons. Notably, ShinyCell2 does not generate a violin plot for the COP population due to the low cell count. Heatmaps are aggregated and lack both single-cell resolution and statistical annotation. (c) Line plot shows memory consumption across datasets ranging from 2,000 to 256,000 cells (with ∼30,000 features per cell).

Second, we evaluated cross-condition gene expression visualization (Fig. 5b) using a published oligodendroglia dataset from healthy and experimental autoimmune encephalomyelitis (EAE) mouse samples [20]. Our goal is to rapidly assess whether the interferon response gene *Stat1* was differentially expressed between disease states across multiple cell types. GUANACO’s split violin plots enabled direct, side-by-side comparisons across both cell types and conditions, with inline statistical testing for significance. This analysis revealed three cell types, astrocytes, mature oligodendrocytes (MOL), and oligodendrocyte precursor cells (OPC), that exhibited significantly elevated *Stat1* expression in EAE samples. Beyond violin plots, GUANACO supports single-cell-resolution heatmaps for visualizing expression patterns across cell types and conditions, with annotation bar providing cell composition information. The relatively small number of COP and MiGl cells, as shown in the heatmap and in Additional file 1: Fig. S1a, may explain the lack of significant group differences for these types, despite apparent differences in their violin plot distributions. Users can zoom in on specific cell types to explore detailed expression profiles across groups (Fig. S1b) or examine overall pseudo-bulk expression using matrix plots (Fig. S1c). In contrast, ShinyCell2 does not support multi-variable faceting and heatmaps at single-cell-resolution. Comparing conditions requires manually switching between separate plots, making visual comparison indirect and potentially less reliable when y-axis ranges differ, as in the violin plots of Fig. 5b. In the case of Vitessce, violin plots are not fully integrated into the selection workflow and were therefore excluded from this comparison.

Finally, we benchmarked the performance between GUANACO and ShinyCell2, as both rely on server-side rendering and offer comparable visualization capabilities. Using datasets ranging from 2,000 to 256,000 cells (with ∼30,000 features each), GUANACO consistently requires ∼50-67% less memory than ShinyCell2’s peak memory during runtime (Fig. 5c). The memory usage was assessed by running both the applications simultaneously and monitoring memory activity from launch until all data and plots were fully loaded, with performance reported as the peak memory usage of each application individually. Notably, neither tool consumes additional memory once the data and plots are loaded. Even then, GUANACO uses ∼50% of ShinyCell2’s peak memory. This difference may be attributed to the coding environments. Shinycell2 runs using R libraries which typically incur higher memory overhead, it also requires converting data from Seurat/SCE objects into HDF5, leading to additional memory consumption. In comparison GUANACO runs using Python directly using backed AnnData objects.

### Flexible and cost-effective deployment with GUANACO

GUANACO has been developed using Python’s Dash platform and Plotly, with the goal of enabling deployment through a single command. Two deployment options are provided: via Docker or directly through the console using the system’s Python environment. Docker typically comes with a preset environment, making it easier to run locally. However, deploying an app with Docker on a server can be somewhat challenging because it requires proper setup to function effectively. Whereas using the system’s Python is a simpler option for both local and server use, particularly for users with limited technical knowledge, as it can be done with a single command. Hence, GUANACO has been designed to make it accessible for users with different levels of Python or Docker expertise.

GUANACO is developed to load data partially in the memory instead of all at once, reducing memory requirements and enabling faster loading of larger datasets with minimal RAM. Hence, for local deployment, a device with 8 GB of RAM is sufficient to handle datasets up to 4 GB in size, including a maximum of 150,000-200,000 cells. For cloud-based deployment, the minimum recommended configuration is 2 CPUs and 4 GB of RAM, which can process datasets up to 500 MB (∼80,000 cells). The storage requirements depend on the dataset size, but generally, 5-10 GB is adequate for storing and deploying GUANACO. Lightweight datasets (<150 MB) can even be hosted on a free AWS EC2 instance. In our current DEMO setup, we deployed GUANACO on an AWS EC2 t4g.small instance (2 CPUs, 2GB RAM), serving a 200 MB PBMC dataset. This instance costs ∼ €6 and operates for 730 hours per month. We have also implemented Nginx as a load balancer to optimize request handling. Full deployment instructions are provided in the Deployment section of the GUANACO User Guide.

GUANACO’s deployment is straightforward because our codebase is compact; a single instance is sufficient for deployment given that no other heavy application is running on that instance. Containerization via Docker is optional and generally recommended for data privacy or reproducibility needs. If server requirements are met, GUANACO can be deployed on a user’s own server ensuring data security. Compared to the ShinyCell2 deployment, GUANACO has a more cost-effective option with longer active hours. For an easy deployment of ShinyCell2, ShinyApp is the best choice, as it can support the essential and resource-intensive packages needed to run ShinyCell2 effectively. However, ShinyApp has limited active hours, and maintaining a web application for our DEMO PBMC dataset over a longer duration, with resource allocation comparable to GUANACO, would cost approximately €38+VAT per month in ShinyApp, while still imposing certain limitations.

## Discussion

Currently, extracting biological insight from single-cell multimodal data is inherently an iterative process. Analysts often move back and forth between code and static plots to refine hypotheses, a “human-in-the-loop” approach critical for improving interpretation and decision-making [21]. However, such workflow is often time consuming and constrained by the limited expressiveness of static visualizations. More systematic integration of interactive visualization into the analytical workflow can accelerate discovery and deepen biological understanding, allowing users to reshape views, pivot perspectives, and uncover complex biological patterns in real time. While current single-cell visualization ecosystems are largely designed for end-stage communication, extending seamless interactivity to the early, exploratory phases of analysis represents an exciting area for the field.

To achieve this, we developed GUANACO, a lightweight platform that reduces the burden of repeated code–plot iterations by enabling interactive exploration through linked plot views and layered cell selection. In practice, GUANACO supports intuitive, data-driven discovery and accommodates common single-cell tasks such as multifactor comparisons, subcluster specific investigations, and conditional filtering.

While providing these capabilities, it is worth noting that interactive visualization is not a panacea. It enhances analysis only when applied purposefully and with awareness of context. Excessive or poorly structured interactions can overwhelm users, introduce noise, and obscure the analytical focus. GUANACO mitigates these risks by employing a progressive disclosure strategy with a layered filtering workflow. Advanced controls are hidden by default and revealed only when needed, helping to maintain a clean and focused interface. Interactive elements are included only when they resolve limitations of static plots, while potentially distracting features are avoided. For example, tooltips are enabled in violin and dot plots where precise values offer interpretive value but are disabled in heatmaps and embeddings where such information is less informative. Filtering is structured in stages: starting from data modality selection, progressing through global cell filtering, free selection in the embedding, and finally plot-specific refinement. This stepwise design allows users to build contextual understanding gradually, without losing track of the data.

There are, however, framework-related limitations. Although Dash/Plotly provides a flexible foundation for matrix and track visualizations and Python simplifies server-side computation, GUANACO is currently limited in coordinating interactions between matrix-based plots and genome tracks because Dash Bio-igv currently does not expose selection events needed for cross-view linking and only supports pre-generated track files rather than tracks generated dynamically from AnnData objects that can be sliced or subset interactively. However, this design by igv.js prioritizes data safety, ensures better storage management, and helps reduce cost. Users can still prepare track files based on metadata of interest and select among predefined sessions to compare pre-defined data views.

The first version of GUANACO primarily focuses on visualizing single-cell RNA-seq and ATAC-seq data. Further work will extend this framework to incorporate additional modalities, such as genomic variants, copy-number profiles, and spatial multi-omics, while preserving low memory overhead alongside the same seamless, interactive workflow. Each of these modalities poses distinct challenges, and their integration will require both new back-end converters and front-end components capable of synchronizing genomic loci and spatial coordinates with the existing matrix track frameworks.

## Conclusions

GUANACO (v1.0.0) is a Dash based Python framework for visualizing matrix and genome track data, enabling the interactive exploration of preprocessed scRNA-seq and scATAC-seq datasets. It provides a unified interface designed for bench scientists with varying levels of computational expertise, offering flexible and intuitive approaches for biological data exploration. We demonstrate that GUANACO supports a broad range of common exploratory tasks in single-cell studies and delivers easier operation and more intuitive visualization compared to existing tools.

Its optimized user experience, low memory footprint, and cost-effective deployment make it straightforward to integrate into existing research workflows for both data exploration and sharing. As an open-source Python tool, GUANACO contributes to the growing ecosystem of reproducible and accessible bioinformatics software. We anticipate that it will help researchers interrogate complex single-cell datasets with greater efficiency and deeper biological insight.

## Methods

### Implementation of GUANACO

GUANACO v1.0.0 is built with Dash, a flexible Python framework for creating interactive web applications, and is designed for visualizing single-cell multi-omics data. The platform integrates matrix-based and coordinate-based genome track views with multiple interactive plot types, and diverse cell and gene selection methods. It also offers specialized features such as statistical testing for group comparisons in violin plots, transcription factor binding site annotation across open chromatin regions, and motif information search. Additionally, GUANACO provides flexible controls and a rich library of colormaps to facilitate the generation of publication-ready figures.

All statistical analyses in GUANACO matrix-based view are implemented using the SciPy [22] and statsmodels [23]. The platform supports four distinct analysis modes: Mode 1 (Single metadata): For comparing expression across groups defined by a single metadata variable. The default method is the non-parametric Mann-Whitney U test for binary comparisons or Kruskal-Wallis test for multiple groups (>2), which assess differences in distributions without assuming normality. Parametric alternatives (t-test for two groups, one-way ANOVA for multiple groups) are available when normality assumptions are met. Mode 2 (Faceted comparisons): For comparing expression between groups of a secondary metadata variable within each level of a primary metadata variable. Statistical tests are performed independently within each facet, using Mann-

Whitney U or Kruskal-Wallis tests by default. Mode 3 (Linear model): Implements a linear regression model (expression ∼ meta1 + meta2) using ordinary least squares to evaluate the main effects of two metadata variables simultaneously. Mode 4 (Mixed effects model): Implements a mixed-effects model (expression ∼ meta1 + (1|meta2)) where meta2 is treated as a random effect. When convergence issues occur, the analysis falls back to a pseudo bulk approach, averaging expression values by meta2 groups before performing t-tests or ANOVA.

The coordinate-based genome track view in GUANACO is adapted from the Integrative Genomics Viewer (IGV), with consistent cell type colors matched to those in the matrix-view visualizations. When zooming into specific peak regions, transcription factor binding motifs from the JASPAR database that overlap with these regions are automatically displayed. Detailed motif information can be searched via the integrated motif search box, implemented using the pyJASPAR API [24], which provides access to the JASPAR 2024 database of transcription factor binding profiles. When a user queries a motif by its ID (e.g., MA1972.1), GUANACO retrieves the corresponding position frequency matrix and generates a sequence logo visualization. The motif visualization employs information content calculated using Shannon entropy. Specifically, the position frequency matrices are first adjusted with pseudo counts, then normalized to calculate position-specific probabilities ( 𝑝_*ai*_ ). The information content at each position is computed as *E_Si_* = Σ𝑝_*ai*_ × *log*_2_(𝑝_*ai*_/*q_a_*), where *q_a_* represents uniform background probabilities (0.25 for each nucleotide). The heights of nucleotides in the sequence logo are determined by multiplying the position-specific probabilities by the information content (heights = 𝑝_*ai*_ × *E_Si_* ), visualized using the logomaker library [25] with a maximum theoretical height of 2 bits.

### Datasets and preprocessing

We used two publicly available PBMC single-cell multiome datasets to demonstrate the interface and visualization capabilities of the GUANACO. The PBMC 3k and 10k datasets were downloaded from the 10x Genomics datasets portal https://www.10xgenomics.com/datasets and processed to produce an .h5mu file containing both RNA and ATAC modalities. Briefly, for each dataset, we read the 10x Multiome matrices into a MuData object, yielding paired scRNA-seq and scATAC-seq layers with feature-interval annotations. For ATAC, we computed QC metrics and applied conservative filters to peaks and cells (retained peaks detected in ≥3 cells; kept cells with 300–25000 detected peaks and total ATAC counts of 1000–50000), then located the fragments file and computed nucleosome signal and TSS enrichment to diagnose low-quality barcodes before the downstream analysis. For RNA, we computed total transcript counts and detected genes per cell, retained cells with adequate library size (total transcript counts >1000), and restricted to barcodes present in both layers to ensure strict RNA–ATAC pairing. We normalized RNA counts to 10000 UMIs per cell, log-transformed, selected highly variable genes, scaled, and computed PCA. Neighborhood graphs were then built with n_neighbors = 30 and n_pcs = 10 for visualization and downstream label transfer. To integrate the PBMC 3k and 10k datasets, we recalculated the shared peak set between the two, merged ATAC profiles on this peak set, removed duplicated feature names, and annotated sample origin (Dataset = “10X3k” or “10X10k”). On the RNA side, we concatenated the 3k and 10k objects using the gene intersection, deduplicated variable names, and dropped analysis-specific columns to maintain a clean joint matrix. Finally, we assembled the paired RNA and ATAC layers into a single MuData object (8677 cells post-filtering), harmonized cell-type labels to a coarse-grained set (B cells, CD4+ T cells, CD8+ T cells, NK, DCs, classical and non-classical monocytes), and wrote both the merged ATAC (.h5ad) and joint RNA+ATAC (.h5mu) files for GUANACO visualization.

ATAC-seq signal tracks were generated as BigWig files for genome browser visualization using CamelGuanACoWig function in the scCAMEL toolkit to export single-cell ATAC-seq signal as one BigWig track per cell type. We read peak-by-cell matrices from AnnData (.h5ad) or MuData (.h5mu; modality “atac”) with anndata and muon (versions recorded in the repository). Peaks were obtained from adata.var[[’Chromosome’,’Start’,’End’]] (or chrom/start/end) when present; otherwise var_names formatted as chrom:start-end were parsed with automatic “chr” prefix enforcement (toggleable). Intervals were filtered to a user-defined chromosome allowlist (human default: chr1–22, X, Y) and sorted by (chrom, start, end); malformed entries (end ≤ start) were removed. Optionally, overlaps were adjusted using PyRanges clustering by shifting each interval’s right boundary to (next_start − 1) and dropping non-positive-length results. Matrices were retained in CSR format and columns re-indexed to the sorted interval order without densification. For aggregation, cells were grouped by adata.obs[celltype_key] (default Cluster) and accessibility was summarized per peak per cell type (default mean; sum available). Chromosome sizes for the BigWig header were provided via a two-column chrom.sizes file or inferred as max(end) per chromosome from the interval set. Final values were optionally scaled (default scale=1.0) and written with pyBigWig as one file per cell type, using chunked inserts for large peak sets. Parameters controlling behavior included allow_chroms, resolve_overlaps, enforce_chr_prefix, aggregation, scale, and outdir; all defaults and exact package versions are specified alongside code in scCAMEL [26] documentation and the repository.

For motif track visualization in Fig. 2, we employed the JASPAR TFBS extraction tool to predict transcription factor binding sites across the PBMC datasets. Peak regions were collected from all cell types and merged to create a union set for binding site analysis. The JASPAR2024 human genome (hg38) database served as the reference for transcription factor binding sites. Without pre-specified transcription factor lists, we applied a stringent score threshold of 400 to filter high-confidence binding sites, consistent with UCSC genome browser standards.

The processed oligodendrocyte scRNA-seq data from healthy and EAE mice were downloaded from the UCSC Cell Browser (https://cells.ucsc.edu/?ds=olg-eae-ms+eae-multiomics+activity) and converted into h5ad files using Scanpy for all visualization comparisons shown in Fig. 5. To facilitate cross-condition analysis, we harmonized cell type annotations by merging molecular subtypes under the ’Celltypes’ classification. Condition-specific cell type labels (such as OPC_Ctr and OPC_EAE) were consolidated into general cell type categories under a new ’Celltypes’ annotation, enabling direct comparison of cell populations between healthy and disease states while preserving the original detailed annotations for subset analyses.

## Declarations

### Ethics approval and consent to participate

The datasets analyzed were previously published and made publicly accessible by the original authors, who confirmed all relevant ethical approvals and participant consents.

## Consent for publication

Not applicable.

## Availability of data and materials

The processed PBMC multi-omics dataset and the oligodendrocyte EAE-MS RNA dataset used in this study are available on Zenodo at https://doi.org/10.5281/zenodo.17151850. A Jupyter notebook illustrating step-by-step conversion to BigWig file is provided here: https://github.com/Systems-Immunometabolism-Lab/guanaco-viz/blob/main/TACoWig_Template.ipynb.

GUANACO is freely available as open-source software under the GNU General Public License version 3. The source code, along with comprehensive documentation, is available on GitHub (https://github.com/Systems-Immunometabolism-Lab/guanaco-viz/) and archived on Zenodo (https://doi.org/10.5281/zenodo.17152010). For ease of deployment, a Docker image is provided at https://hub.docker.com/r/systemsimmunometabolismlab/guanaco-viz/. An interactive demonstration showcasing GUANACO’s capabilities is accessible at https://guanaco-demo.chen-sysimeta-lab.com/.

## Authors’ contributions

X.Z., Y.H., and P.C. conceived and designed the project. P.C. managed and supervised the project. X.Z. and Y.H. acquired and processed the data. X.Z. and Q.X. developed the GUANACO. M.K. implemented the deployment framework. X.Z. and M.K. conducted case studies and performed comparisons with other tools. X.Z., M.K., Y.H., and P.C. wrote the manuscript. All authors reviewed and approved the final manuscript.

## Acknowledgements

We are grateful to Artur Khaziev for creating the GUANACO logo and comments on the user guide and GUANACO interface design. We wish to acknowledge the CSC-IT Center for Science (Finland) for computational resources and the open access funding provided by Karolinska Institutet. We thank Gonçalo Castelo-Branco and Eneritz Agirre (Department of Medical Biochemistry and Biophysics, Karolinska Institutet) for coordinating the oligodendroglia dataset.

## Supplementary Information

Additional file 1: Supplementary figures

Additional file 2: Comparison of GUANACO with existing visualization tools

## References

1. Lim J, Park C, Kim M, Kim H, Kim J, Lee D-S, Lim J, Park C, Kim M, Kim H, et al: Advances in single-cell omics and multiomics for high-resolution molecular profiling. Experimental & Molecular Medicine 2024 56:3 2024-03-05, 56.

2. Lähnemann D, Köster J, Szczurek E, McCarthy DJ, Hicks SC, Robinson MD, Vallejos CA, Campbell KR, Beerenwinkel N, Mahfouz A, et al: Eleven grand challenges in single-cell data science. Genome Biology 2020 21:1 2020-02-07, 21.

3. Virshup I, Rybakov S, Theis FJ, Angerer P, Wolf FA: anndata: Access and store annotated data matrices. Journal of Open Source Software 2024/09/16, 9.

4. Butler A, Hoffman P, Smibert P, Papalexi E, Satija R, Butler A, Hoffman P, Smibert P, Papalexi E, Satija R: Integrating single-cell transcriptomic data across different conditions, technologies, and species. Nature Biotechnology 2018 36:5 2018-04-02, 36.

5. MD L, FJ T: Current best practices in single-cell RNA-seq analysis: a tutorial - PubMed. Molecular systems biology 06/19/2019, 15.

6. Blanco-Carmona E: Generating publication ready visualizations for Single Cell transcriptomics using SCpubr. bioRxiv 2022-03-01.

7. Program CCS, Abdulla S, Aevermann B, Assis P, Badajoz S, Bell SM, Bezzi E, Cakir B, Chaffer J, Chambers S, et al: CZ CELLxGENE Discover: a single-cell data platform for scalable exploration, analysis and modeling of aggregated data. Nucleic Acids Research 2025/01/06, 53.

8. Speir ML, Bhaduri A, Markov NS, Moreno P, Nowakowski TJ, Papatheodorou I, Pollen AA, Raney BJ, Seninge L, Kent WJ, Haeussler M: UCSC Cell Browser: visualize your single-cell data. Bioinformatics 2021 Jul 9, 37.

9. Keller MS, Gold I, McCallum C, Manz T, Kharchenko PV, Gehlenborg N, Keller MS, Gold I, McCallum C, Manz T, et al: Vitessce: integrative visualization of multimodal and spatially resolved single-cell data. Nature Methods 2024 22:1 2024-09-27, 22.

10. Chen BJ, Lim YY, Yang X, Wang L, Rackham OJL, Ouyang JF: ShinyCell2: An extended library for simple and sharable visualisation of spatial, peak-based and multi-omic single-cell data. bioRxiv 2025-04-23.

11. Bredikhin D, Kats I, Stegle O, Bredikhin D, Kats I, Stegle O: MUON: multimodal omics analysis framework. Genome Biology 2021 23:1 2022-02-01, 23.

12. Wolf FA, Angerer P, Theis FJ, Wolf FA, Angerer P, Theis FJ: SCANPY: large-scale single-cell gene expression data analysis. Genome Biology 2018 19:1 2018-02-06, 19.

13. RP D, BL S, MB K, D J, H T, C B, DA H, KJ H: Regulation of the interleukin-7 receptor alpha promoter by the Ets transcription factors PU.1 and GA-binding protein in developing B cells - PubMed. The Journal of biological chemistry 05/11/2007, 282.

14. Robinson JT, Thorvaldsdottir H, Turner D, Mesirov JP: igv.js: an embeddable JavaScript implementation of the Integrative Genomics Viewer (IGV). Bioinformatics 2023/01/01, 39.

15. Rauluseviciute I, Riudavets-Puig R, Blanc-Mathieu R, Castro-Mondragon Jaime A, Ferenc K, Kumar V, Lemma RB, Lucas J, Chèneby J, Baranasic D, et al: JASPAR 2024: 20th anniversary of the open-access database of transcription factor binding profiles. Nucleic Acids Research 2024/01/05, 52.

16. Wong B: Points of view: Color blindness. Nature Methods 2011 8:6 2011-05-27, 8.

17. Okabe M, Ito K: Color Universal Design (CUD): How to make figures and presentations that are friendly to colorblind people. 2002.

18. Coblis — Color Blindness Simulator [https://www.color-blindness.com/coblis-color-blindness-simulator/]

19. Ng SS, De Labastida Rivera F, Yan J, Corvino D, Das I, Zhang P, Kuns R, Chauhan SB, Hou J, Li X-Y, et al: The NK cell granule protein NKG7 regulates cytotoxic granule exocytosis and inflammation. Nature Immunology 2020 21:10 2020-08-24, 21.

20. Meijer M, Agirre E, Kabbe M, Tuijn CAv, Heskol A, Zheng C, Falcão AM, Bartosovic M, Kirby L, Calini D, et al: Epigenomic priming of immune genes implicates oligodendroglia in multiple sclerosis susceptibility. Neuron 2022/04/06, 110.

21. The Interactive Visualization Gap in Initial Exploratory Data Analysis. IEEE Transactions on Visualization and Computer Graphics 2018, 24.

22. Virtanen P, Gommers R, Oliphant TE, Haberland M, Reddy T, Cournapeau D, Burovski E, Peterson P, Weckesser W, Bright J, et al: SciPy 1.0: fundamental algorithms for scientific computing in Python. Nature Methods 2020 17:3 2020-02-03, 17.

23. Seabold S, Perktold J: Statsmodels: Econometric and Statistical Modeling with Python - SciPy Proceedings. *scipy* 2010/5/1.

24. Khan A: pyJASPAR: a Pythonic interface to JASPAR transcription factor motifs.

25. Tareen A, Kinney JB: Logomaker: beautiful sequence logos in Python. Bioinformatics 2020/04/01, 36.

26. Xu Y, Lou D, Chen P, Li G, Usoskin D, Pan J, Li F, Huang S, Hess C, Tang R, et al: Single-cell MultiOmics and spatial transcriptomics demonstrate neuroblastoma developmental plasticity. Developmental Cell 2025, 0.

