## Supplemental Figure 1 for "GUANACO: A Unified Web-Based Platform for Single-Cell Multi-Omics Data Visualization"

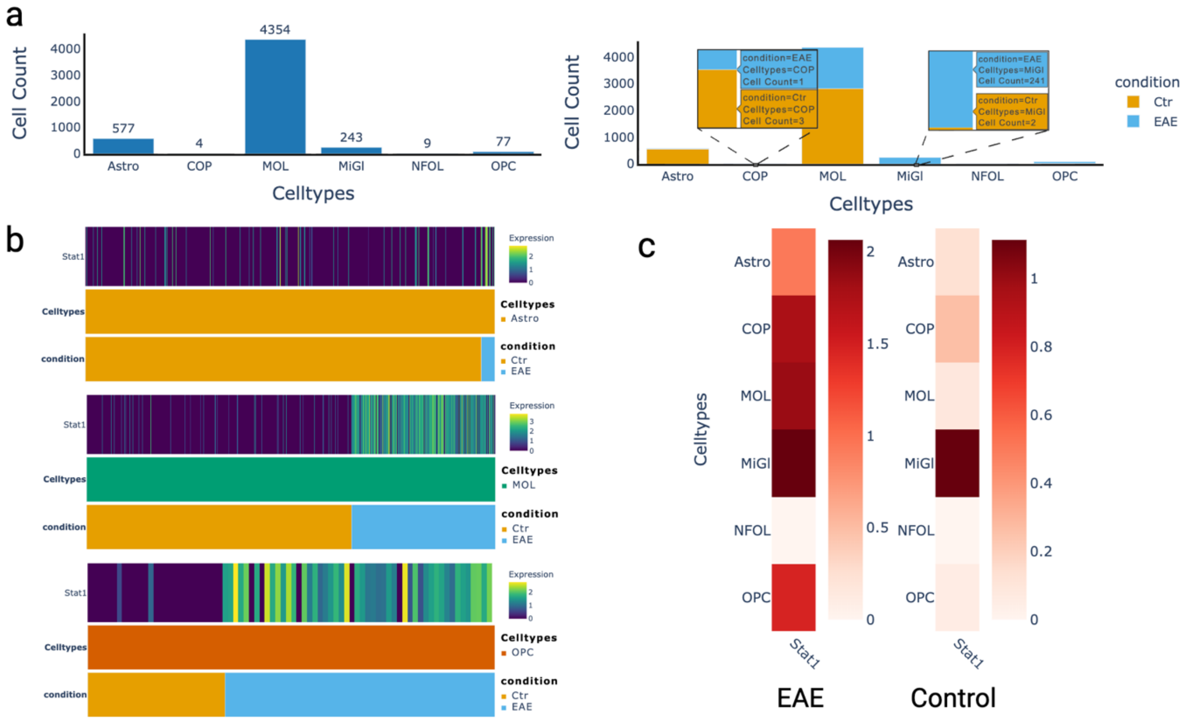


**Figure S1. *Stat1* expression visualized by GUANACO.** a. Bar plots displaying cell counts for each cell type. The left panel shows total cell numbers across all conditions, while the right panel displays counts split by condition. Note that COP and NFOL have low cell numbers, and MiGl shows an imbalanced distribution (predominantly in the Ctrl group). These differences in cell number and distribution help explain the weaker statistical significance observed in some comparisons. b. Single-cell heatmaps showing *Stat1* expression in Astro, MOL, and OPC cells separately, showing higher *Stat1* expression in the EAE group across all three cell types. c. Pseudo-bulk level heatmaps illustrating *Stat1* expression in the EAE and control groups for each cell type.
